# Differential Sensitivity of HSV-1 and PRV to IFN-λ Reveals a Neuron-Specific Antiviral Role for RSAD2

**DOI:** 10.64898/2026.06.01.729199

**Authors:** Stephanie Salazar, Khanh T. Y. Luong, Audrey D. Loaiza, Dillon Cheng, Nir Drayman, Orkide O. Koyuncu

## Abstract

Alpha herpesviruses (α-HV) initially infect mucosal epithelial cells and subsequently establish lifelong latency in the peripheral nervous system (PNS). Herpes simplex virus-1 (HSV-1), a human pathogen persisting in the majority of the adult population, shares neuroinvasive properties with Pseudorabies virus (PRV), a swine α-HV, commonly used as a model α-HV. Utilizing primary peripheral neuronal cultures, we previously showed that IFN-λ pre-treatment significantly reduced PRV yield. In this paper, we further characterized the early and late neuronal responses to IFN-λ by RNA-seq, and the antiviral potential of this response against HSV-1. Notably, HSV-1 exhibited neuron-specific resistance to IFN-λ mediated antiviral responses both in murine primary neurons and human neuronal cells. An ICP34.5-deficient HSV-1 (Δ34.5) mutant showed IFN-λ sensitivity in neurons, while replicating normally in untreated neurons showing that ICP34.5 is responsible for the neuron specific IFN-λ resistance of HSV-1. Our results further demonstrate that RSAD2 is strongly induced by IFN-λ in neurons, localizing to ER-associated membranes, and effectively restricting α-HV protein synthesis in the absence of ICP34.5. siRNA-mediated RSAD2 knockdown in IFN-λ-primed primary neurons largely restored replication of Δ34.5 HSV-1, highlighting the role of this IFN-λ induced host factor in neuronal infections. Together, neuronal IFN-λ-induced RSAD2 and HSV-1 ICP34.5 define a neuron-specific antagonistic mechanism that collectively determines the replication efficiency of HSV-1 in the PNS.

## Introduction

Most α-HV infections begin in the periphery at the mucosal epithelium, such as the oral, genital, nasal, or oropharyngeal mucosa [1]. α-HV infections within the mucosal epithelia result in productive replication, yielding hundreds to thousands of progeny per infected cell [2]. Some of these newly made virus particles spread the infection to neighboring epithelial cells, fibroblasts, or nerve endings. Despite recent discoveries, there is a large knowledge gap between the connection of the mucosal epithelia and the PNS junction.

Mucosal epithelial cells are a subset of epithelial cells that line the mucosal barriers of the genital, respiratory, and gastrointestinal systems. The innate immune responses of mucosal epithelia serve as a critical barrier, preventing virions from reaching peripheral neurons during primary infection. Epithelial interferons (IFNs) play a key role in orchestrating these antiviral responses. Type I IFNs (IFN-α/β) and type III IFNs (IFN-λ) mediate intrinsic and natural antiviral immunity in the infected mucosal epithelia [3, 4]. Unlike type I IFN receptors (IFNAR1/2), type III IFN receptors, composed of IFNLR1/IL-10RB heterodimer, are not ubiquitously expressed and are instead preferentially expressed on mucosal epithelial cells and immune cells, such as neutrophils, dendritic cells, and B cells [3, 5]. Interestingly, IFN-λ production increases with the cell polarity, and the quality of the IFN response is determined by the differentiation state of mucosal epithelial cells [4]. Functional studies have demonstrated that type III IFN treatment of mucosal epithelial cells upon α-HV infection had a higher protective response when compared to type I IFN treatment [3, 6, 7]. Importantly, intravaginal pre-treatment of IFN-λ prior to HSV-2 infection in a murine model elicited a potent antiviral response [8]. Despite signaling through distinct receptors, both type I and type III IFNs activate overlapping downstream pathways (e.g., JAK-STAT pathway) involving signal transducer and activator of transcription (STAT) phosphorylation and IFN-stimulated gene (ISG) induction.

Given that α-HV infection begins at the mucosal epithelium and that peripheral nerve endings are the first neuronal components exposed to this cytokine-rich microenvironment, we hypothesized that neurons respond to type III IFN signaling and establish an antiviral state. We have previously shown that peripheral neurons respond to IFN-λ through a noncanonical pathway. Unlike fibroblasts, which activate both STAT1 and STAT2 upon IFN-λ treatment, neurons exhibit selective STAT1 phosphorylation without STAT2 activation leading to the expression of a subset of ISGs [9]. In this study, we further characterized this response in primary neurons through RNA-seq and found that the STAT1-dominant signaling drives a biphasic transcriptional response: early neuronal responses included genes involved in neuronal structure, signaling, and metabolic adaptation, followed by delayed but robust induction of antiviral ISGs. Compared with type I IFN, which elicits rapid, high-magnitude ISG induction, IFN-λ produced a delayed and lower magnitude response in neurons.

Among the ISGs, RSAD2 (radical S-adenosyl methionine domain-containing 2), also known as viperin, stood out as the most upregulated gene after 24 h of IFN-λ treatment in primary neurons. RSAD2 is known for its broad-spectrum antiviral activity and a unique enzymatic mode of action [10]. Structurally, RSAD2 contains a radical S-adenosyl-L-methionine (SAM) enzyme domain with a 4Fe-4S cluster, a cofactor typically associated with metabolic enzymes rather than classic immune effectors [11, 12]. RSAD2 catalyzes the conversion of cytidine triphosphate (CTP), a canonical nucleotide, to synthesize 3’-deoxy-3’,4’-didehydro-CTP (ddhCTP), an antiviral ribonucleotide analog that terminates viral RNA synthesis [9, 13, 14]. This chain-terminating analog inhibits viral RNA-dependent RNA polymerases, effectively blocking the elongation of nascent viral RNA chains [15].

We further investigated the antiviral potential of the neuronal IFN-λ response and RSAD2 induction against HSV-1 infection. While our previous findings showed that PRV was highly sensitive to IFN-λ-mediated restriction in neurons, HSV-1 replicated efficiently despite effective STAT1 phosphorylation and ISG expression. Remarkably, this neuron-specific resistance is conserved across species as both rat superior cervical ganglionic neurons (SCG) and human SK-N-SH neuroblastoma cells failed to restrict HSV-1 infection, ruling out species-specific differences. These findings indicate that HSV-1 has evolved specialized mechanisms to overcome antiviral defenses particularly in neurons.

HSV-1 encodes unique neurovirulence factors that are absent in many other herpesviruses. Most notably, ICP34.5 antagonizes host antiviral responses through multiple complementary mechanisms [16, 17]. ICP34.5 recruits protein phosphatase 1α (PP1α) to dephosphorylate elongation initiation factor 2 alpha (eIF2α) and reverses translational shutoff, binds Beclin-1 to inhibit autophagy, and inhibits TBK1-mediated IRF3 phosphorylation to attenuate IFN production [18–20]. Similar to PRV, an HSV-1 mutant lacking ICP34.5 was sensitive to the neuronal IFN-λ response. In primary rat neurons and a human neuronal cell line that were IFN-λ pretreated and infected with ICP34.5-deficient HSV-1 (Δ34.5), RSAD2 expression remained high, its punctate perinuclear localization was preserved, eIF2α phosphorylation persisted, and viral protein synthesis was severely impaired. This infection-limiting effect was mostly restored during Δ34.5 infection when RSAD2 expression was downregulated by RNAi in IFN-λ treated primary neurons.

Collectively, this study emphasizes the role of IFN-λ in activating a potent antiviral response with rapid, high-magnitude restriction in fibroblasts and slower, but sustained responses in PNS neurons. However, these defenses are not equally effective against neuroinvasive α-HV infections. While a varicellovirus, PRV remains vulnerable to neuronal IFN-λ response, simplexviruses (e.g. HSV-1) encoding ICP34.5, have evolved specialized countermeasures that neutralize neuron-specific antiviral mechanisms. Such evolutionary divergence may explain the widespread HSV-1 infections and their capacity to establish lifelong latency in the nervous system. This work identifies the IFN-λ-RSAD2-eIF2α axis as a neuronal antiviral defense limiting viral protein synthesis and ICP34.5 as an HSV-1 factor counteracting this shut off not only by targeting eIF2α, but RSAD2 as well.

## Results

### Temporal ISG induction in IFN-λ-treated primary neurons

Our previous data showed the selective activation of STAT1, but not STAT2 in peripheral neurons, suggesting that type III IFNs might activate a noncanonical antiviral response different from the well-known ISGF3 (IFN-stimulated gene factor 3) response. To gain an unbiased view of this alternative signaling pathway, we conducted RNA sequencing to map the transcriptional landscape of IFN-λ-treated peripheral neurons. We performed RNA-sequencing at early (3 h) and late (24 h) time points after primary superior cervical ganglionic (SCG) neurons were treated with IFN-λ2. At 3 h post-treatment, we observed distinct changes in gene expression. Notably, the most significantly downregulated genes included H19 (a long non-coding RNA implicated in growth control and neural development), Nos1 (a neuronal nitric oxide synthase involved in neurotransmission), and Dcx (a marker of immature neurons involved in neuronal migration) (Figure 1A) [21, 22]. Conversely, Ahnak, a gene involved with membrane repair and calcium signaling, was among the most significantly upregulated (Figure 1A) [23]. Gene ontology analysis using *gProfiler* further confirmed these enrichments. By 24 hours post-treatment, a shift toward ISG induction was observed, with 32.6% of the significantly induced genes being associated with innate immune response pathways (Fig S1). RSAD2 (viperin) emerged as the most robustly upregulated transcript at this later stage (Figure 1B). Other top hits in this later stage included *Mx1* (a GTPase that inhibits viral replication) and *Usp18* (an ISG15 de-conjugating enzyme that acts as a negative regulator of IFN signaling), classic ISGs that are key in facilitating the antiviral response induced by type I and type III IFNs [24, 25].

**Figure 1.**
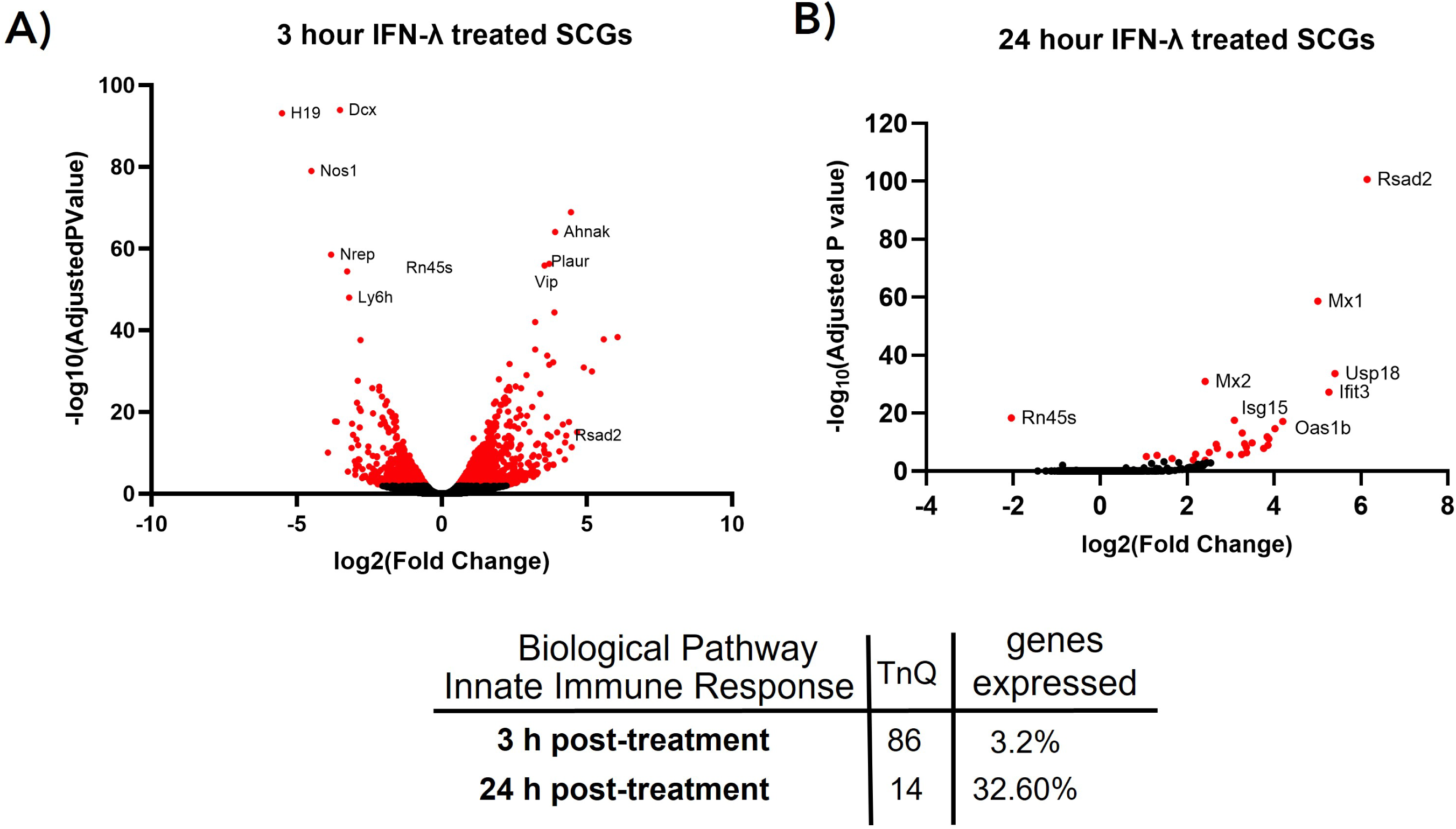
Delayed ISG activation characterizes the IFN-λ response in peripheral neurons. SCGS were treated with rat IFN-λ2 (200 ng/ml) or vehicle for (**A**) 3 h and (**B**) 24 h. Volcano plots represent differential gene expression normalized to GAPDH. Scattered points represent genes; the x-axis is the log 2-fold change, whereas the y-axis is the statistical significance in differential expression. Red dots highlight genes significantly over-or under-expressed after IFN-λ exposure. The abundance of genes that play a role in innate immune responses are determined by using *gProfiler*.

### Dynamic induction of interferon-stimulated genes (ISGs) in IFN-λ- and IFN-β-treated SCGs

To further validate our RNA-sequencing results, we quantified the time-dependent induction of ISGs in primary SCG neurons following treatment with either rat IFN-λ2 (200 ng/ml) or IFN-β (100 U/ml). Normalized gene expression analysis revealed that IFN-λ induced a modest, albeit discernible, increase in ISG transcripts at 3 h post-treatment, which became more pronounced by 8 h and 24 h (Figure 2A). RSAD2 expression in SCG neurons increased in a time-dependent manner following IFN-λ treatment, with modest induction at 3 h, a significant increase by 8 h, and the highest expression at 24 h (p < 0.05 and p < 0.05 relative to 3 h, respectively; Figure 2A). In contrast, canonical ISGs; OAS1a and IFIT1 exhibited minimal induction following IFN-λ treatment and did not significantly increase over time.

**Figure 2.**
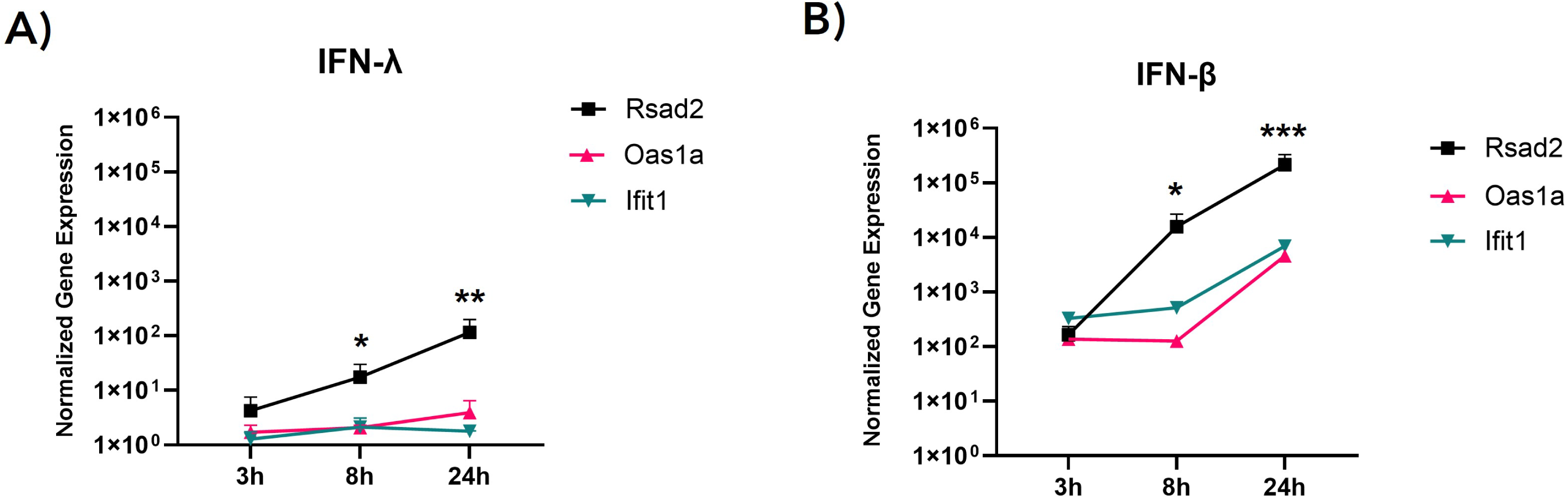
Time-course qPCR analysis of ISG induction in IFN-λ-treated SCGs. Normalized gene expression (2^−ddCt^) in superior cervical ganglion (SCG) neurons following (**A**) rat IFN-λ2 treatment (200 ng/ml) or (**B**) rat IFN-β treatment (100 U/ml) at 3 h-, 8 h- 24 h-post treatment. GAPDH was used as a housekeeping gene. Untreated SCGs served as the control. Statistical significance analyzed via Student’s *t* test (n ≥ 3).

In comparison, IFN-β induced a higher and earlier expression of ISG transcripts at 3 hours compared to neurons treated with IFN-λ2. IFN-β treatment elicited a rapid and robust ISG response, characterized by a significant increase in RSAD2 expression as early as 8 h (p < 0.05) and further enhancement at 24 h (p < 0.005; Figure 2.B). OAS1a and IFIT1 were also significantly induced by IFN-β at 24 h compared to 3 h (p < 0.05). Importantly, RSAD2 levels were 2.14 × 10^5^ (± 19.6 SEM)-fold higher at 24 h following IFN-β treatment, whereas neurons treated with IFN-λ elicited a 1.16 × 10^2^ (± 83 SEM)-fold increase (Figure 2B). Notably, RSAD2 levels induced by IFN-λ at 24 h were comparable to those induced by IFN-β at 3 h, indicating delayed but sustained ISG activation downstream of IFN-λ signaling.

### HSV-1 mitigates the IFN-λ-induced antiviral response in neurons

To determine whether this bi-phasic IFN-λ neuronal response was comparably effective against HSV-1 infection as we reported against PRV infection, we treated SCG neurons with IFN-λ for 24 h before virus infection. For these experiments, we used a recombinant HSV-1 strain 17 expressing mRFP-VP26 (OK14) [26, 27]. Surprisingly, we did not observe a significant reduction in OK14 viral yield at 48 hours post-infection (hpi) when SCG neurons were pretreated with 200 ng/ml of IFN-λ2 (Figure 3A). In contrast, Rat2 fibroblasts pretreated with IFN-λ exhibited a significant reduction (2.1 × 10² ± 70 SEM-fold) in OK-14 titers after 24 hpi (Figure 3C).

**Figure 3.**
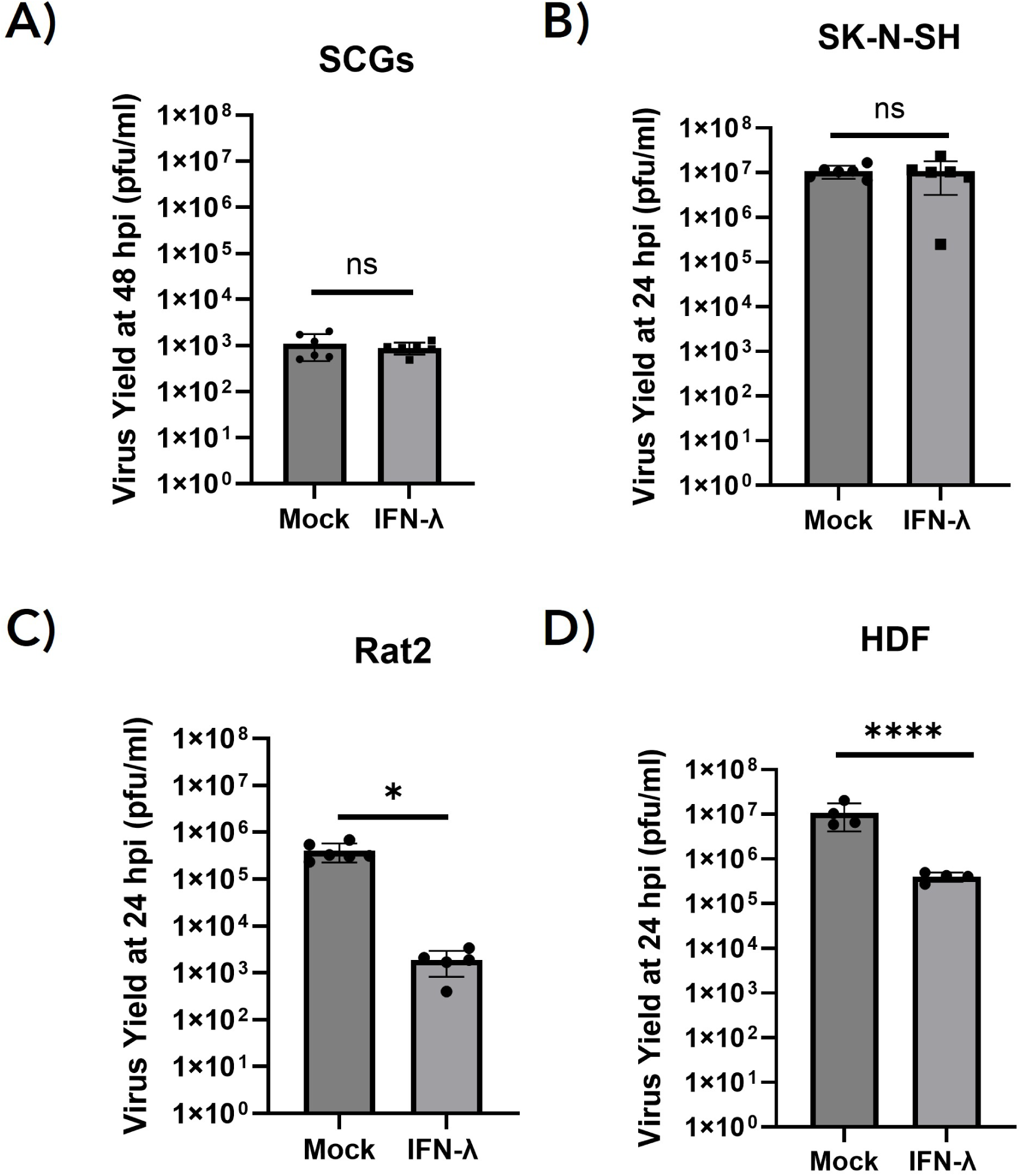
Differential antiviral activity of IFN-λ across neuronal and non-neuronal cell types following HSV-1 infection. (**A**) SCGs or (**C**) Rat2 cells were pre-treated with rat IFN-λ2 (200 ng/mL) for 24 h prior to infection with OK-14 (MOI 1). (**B**) SK-N-SH cells or (**D**) HDFs were pre-treated with human IFN-λ2 (200 ng/mL) for 24 h prior to infection with OK-14 (MOI 1). Virus yield was determined as plaque forming units (pfu). Statistical significance analyzed via Student’s *t* test (n ≥ 3) is indicated as follows: ns, not significant; *, *p* < 0.05, ****, *p* < 0.00005.

SK-N-SH cells (a human neuroblastoma-derived cell line) were used to assess species-specific differences in neuronal IFN-λ responsiveness [28]. This comparison was crucial as human and rat neurons may differ in interferon receptor expression and downstream signaling pathways. SK-N-SH cells pretreated with human IFN-λ2 showed comparable ISG expression (Figure S2A) to SCGs, but no significant suppression of HSV-1 replication was detected (Figure 3B). Interestingly, similar to rat fibroblasts, human dermal fibroblasts (HDFs) primed with human IFN-λ2, exhibited a strong reduction in the virus yield (2.69 × 10¹ (± 8.90 SEM)-fold) (Figure 3D). Overall, these results suggest that IFN-λ induced antiviral activity against HSV-1 is potent in fibroblasts but not in neurons.

### ICP34.5 suppresses IFN-λ-induced antiviral response during HSV-1 infection

Our results show that HSV-1 effectively suppresses the IFN-λ-mediated antiviral responses in neurons, while PRV remains susceptible to it. Although PRV and HSV-1 share a highly conserved genomic structure and encode many homologous genes, there are differences in key genomic regions. Specifically, the inverted repeat (IR) sequences, which are flanked by the unique long (UL) and unique short (US) regions of the viral genome, contain genes that regulate viral pathogenesis and immune evasion (Figure 4A). PRV contains IRs that are relatively compact compared to HSV-1 and lacks certain immunomodulatory genes present in HSV-1, such as ICP34.5, a multifaceted protein essential for mitigating immune responses and counteracting host antiviral pathways. Based on this difference, we hypothesized that the absence of ICP34.5 would make HSV-1 more vulnerable to IFN-λ-induced antiviral activity. To test this, we assessed the replication of an HSV-1 mutant lacking ICP34.5 (Δ34.5) and compared it to its revertant, 34.5R (Figure 4A) [29].

**Figure 4.**
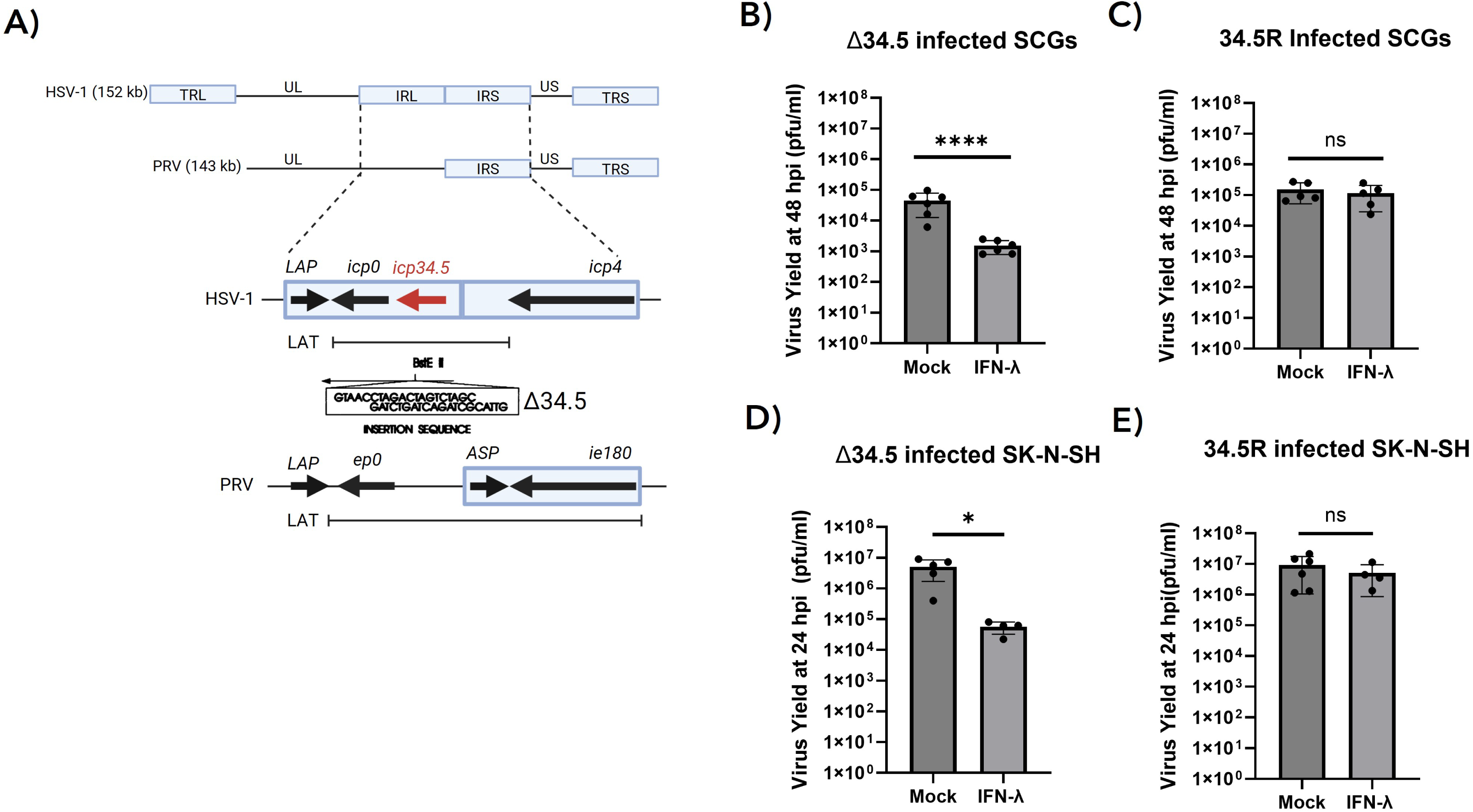
Loss of ICP34.5 renders HSV-1 more susceptible to IFN-λ-mediated restriction in neuronal cells. (**A**) Schematic comparison of the HSV-1 and PRV genomes, highlighting ICP34.5 that is missing in the PRV genome. HSV-1 mutant, Δ34.5 (originally named 17termA) illustration is adapted from Bolovan, et al.[51]. SCGs were pre-treated with rat IFN-λ2 (200 ng/mL) for 24 h prior to infection with (**B**) Δ34.5 or (**C**) 34.5R for 48 h. SK-N-SH cells were pre-treated with human IFN-λ2 (200 ng/mL) for 24 h prior to infection with **(D)** Δ34.5 or (**E**) 34.5R for 24 h. Statistical significance analyzed via Student’s *t* test (n ≥ 4) is indicated as follows: ns, not significant; *, *p* < 0.05, ****, *p* < 0.00005.

In SCG neurons pretreated with rat IFN-λ, the replication of Δ34.5 was significantly reduced compared to untreated controls (p < 0.05; Figure 4B). This indicates that ICP34.5 is essential for HSV-1 to evade the IFN-λ-mediated antiviral response in neurons effectively. Notably, in the absence of IFN-λ, Δ34.5 replicated to titers comparable to the revertant virus (34.5R), demonstrating that the mutant is not intrinsically impaired in replication in SCG neurons. (Figure 4C). Similarly, in human SK-N-SH cells, pretreatment with IFN-λ also resulted in a significant decrease in Δ34.5 titers (Figure 4D), confirming the neuron-specific, rather than species-specific resistance of HSV-1. Likewise, in the absence of IFN-λ, Δ34.5 replicated to similar titers as 34.5R-infected SK-N-SH cells (Figure 4D and 4E). Overall, these findings suggest that the differential sensitivity of HSV-1 and PRV to IFN-λ is linked to the presence of ICP34.5, which enables HSV-1 to suppress host antiviral defenses in neuronal cells.

### Neuronal IFN- λ response does not affect viral gene transcription but reduces viral protein synthesis

To assess the effect of IFN-λ on viral gene expression, we pretreated SCG neurons with IFN-λ before infecting them with 34.5R or Δ34.5 at an MOI of 1. Viral transcripts of immediate-early (ICP27), early (ICP8), and late (gC) genes were measured by Q-PCR at 3 h, 8 h, and 24 h after infection and normalized to GAPDH. Surprisingly, no defect or delay in viral gene expression was detected at any of the examined time points for any of the viral genes analyzed (Figure 5A, B and C). This finding suggests that IFN-λ treatment does not directly impair viral gene transcription in the absence of ICP34.5.

**Figure 5.**
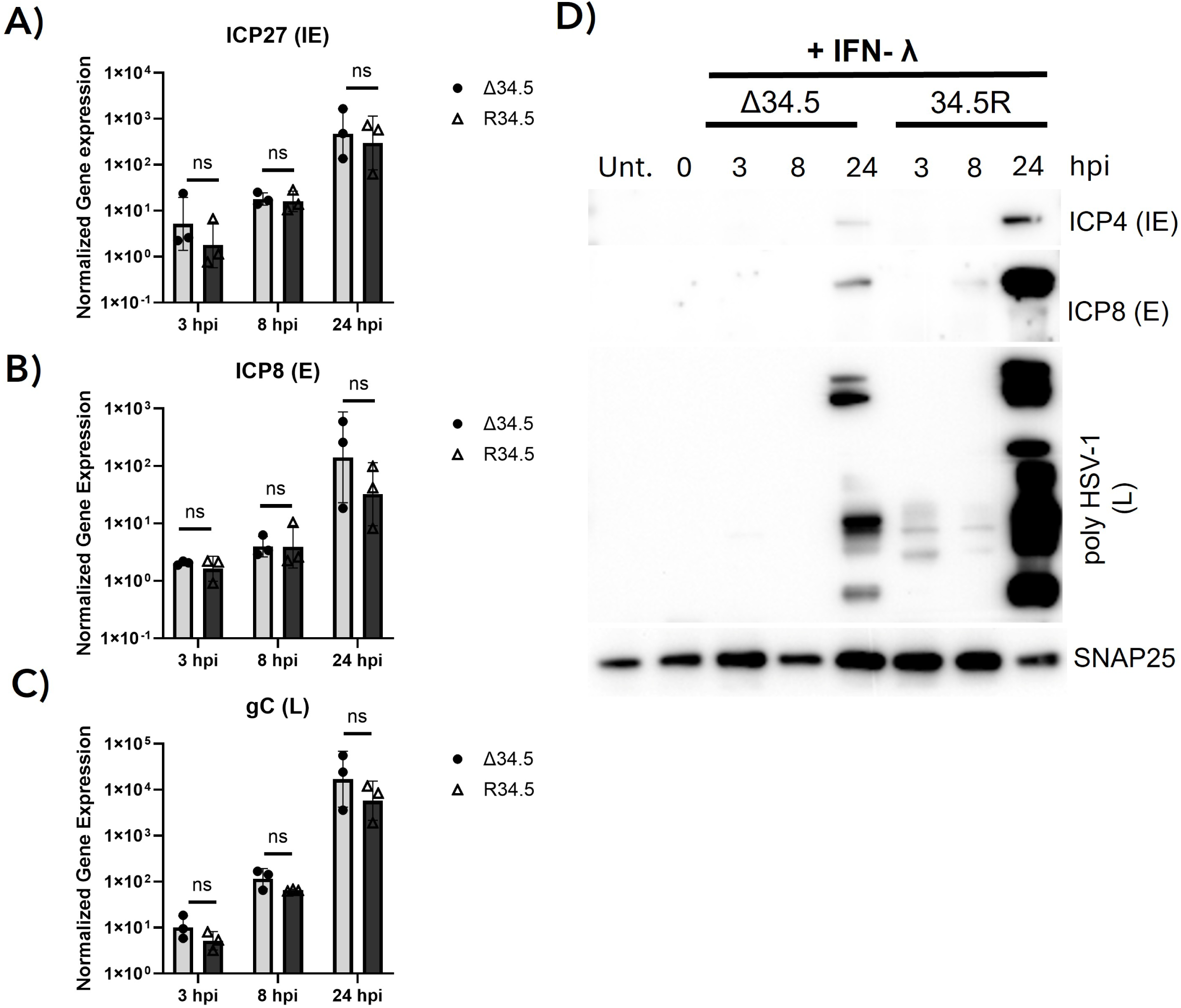
Neuronal IFN-λ response does not alter HSV-1 transcription but interferes with viral protein synthesis when ICP34.5 is absent. qPCR quantification of viral immediate-early (ICP27) (**A**), early (ICP8) (**B**), and late (gC) (**C**) gene expression and protein accumulation (**D**) in SCGs pre-treated with rat IFN-λ2 (200 ng/ml) for 24 h and infected with Δ34.5 or 34.5R at MOI 1 for either 3 h-, 8 h-, or 24 hpi. Statistical significance was performed using Student’s *t* test: *, *p* < 0.05 (n=3 replicates). Immunoblotting was done using antibodies against ICP4, ICP8, poly-HSV-1, and SNAP25.

To determine whether IFN-λ differentially affects viral protein synthesis in the absence of ICP34.5, immunoblot analysis was performed using antibodies against ICP4 (IE), ICP8 (E), and poly HSV-1 (L) in a time-course of (3 h, 8 h, and 24 h) infection. In IFN-λ pre-treated neurons, Δ34.5 infection demonstrated reduced IE, E, and L protein levels compared to 34.5R-infected neurons (Figure 5D). Despite efficient transcription, late protein expression was reduced in SCGs infected with Δ34.5, suggesting that ICP34.5 is essential for sustaining temporally regulated protein synthesis under IFN-λ pressure.

### IFN-λ triggers strong neuronal RSAD2 induction and perinuclear ER localization

Since RSAD2 was detected as the highest induced ISG in the RNA-seq analysis, we further evaluated RSAD2 protein expression and subcellular localization patterns following IFN-λ treatment in primary neurons and human neuronal cell lines. In SCG neurons exposed to IFN-λ2, immunofluorescence staining revealed distinct RSAD2 puncta localized predominantly in the perinuclear region, largely overlapping with the ER marker, calnexin, with minimal presence in untreated controls (Figure 6A). Western blot analysis corroborated these findings, indicating a robust induction of RSAD2 protein at 24 hours post-treatment (Figure 6B).

**Figure 6.**
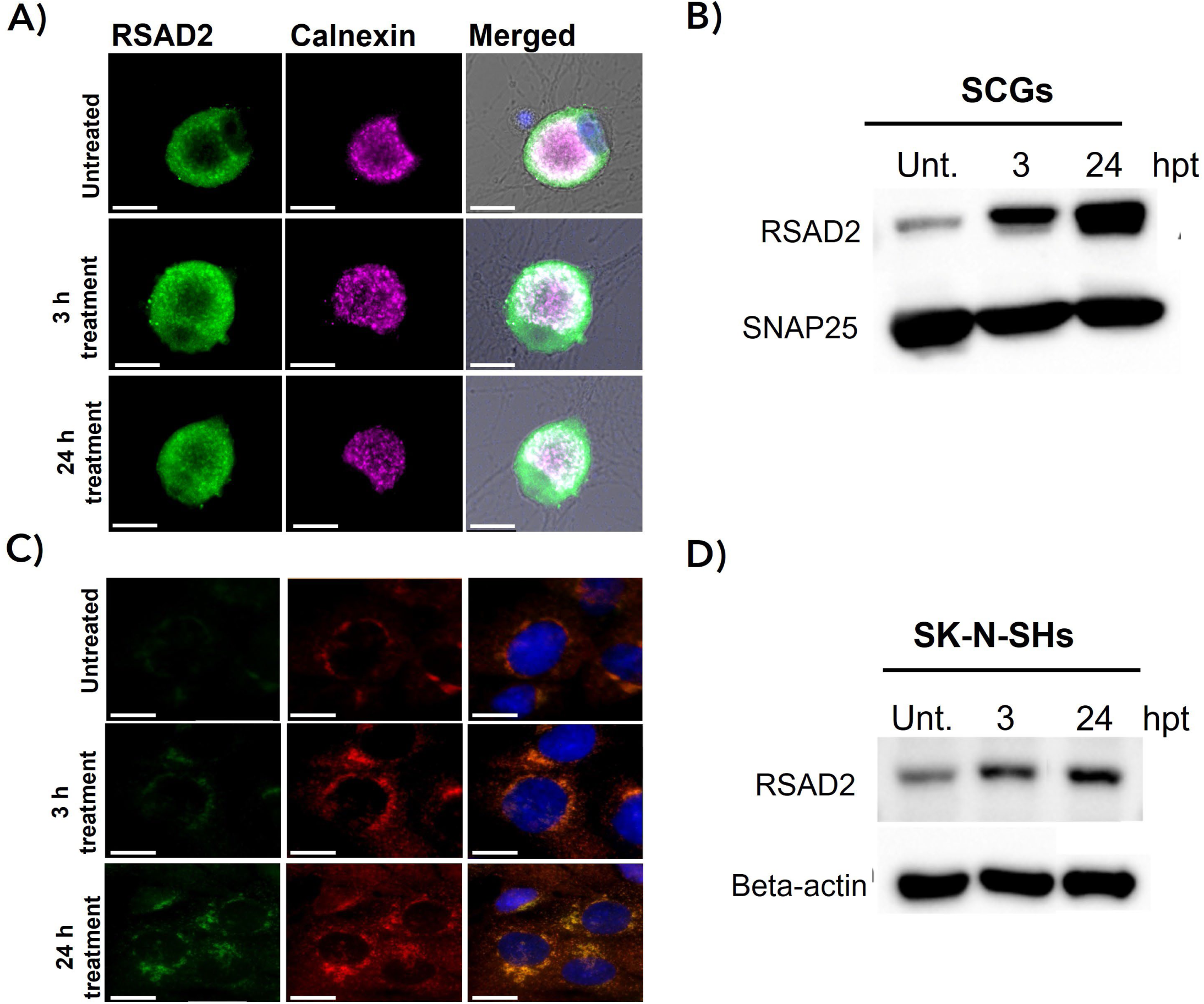
RSAD2 expression increases in SCGs after IFN-λ treatment. (**A**) Immunofluorescence staining of RSAD2 in SCGs after 3 h- and 24 h-post-treatment with rat IFN-λ2 (scale bars, 10 µm). (**B**) Immunoblotting of RSAD2 in SCGs treated with rat IFN-λ2 for 3 or 24 hours. SNAP25 was used as a loading control. (**C**) Immunofluorescence staining of RSAD2 in SK-N-SH cells treated with human IFN-λ2 for 3 or 24 hours (scale bars, 20 μm). (**D**) Immunoblotting of RSAD2 in SK-N-SH cells treated with human IFN-λ2 for either 3 or 24 hours. Beta-actin was used as a loading control.

A comparable temporal pattern was observed in human SK-N-SH cells, where immunofluorescence analysis revealed perinuclear RSAD2 puncta as early as 3 hours, with a pronounced increase by 24 h colocalizing to ER (Figure 6C). Notably, in both SCGs and SK-N-SH cells, RSAD2 perinuclear localization remained sustained across all examined time points, indicating that this spatial pattern is a stable and conserved feature of IFN-λ signaling in neurons. Immunoblotting confirmed upregulation of RSAD2 protein at both time points, 3 h and 24 h post-treatment (Figure 6D). These results collectively demonstrate that IFN-λ significantly upregulated RSAD2 protein expression in primary neurons and neuronal cells.

### HSV-1 infection alters RSAD2 expression through ICP34.5 protein to ensure efficient viral protein synthesis in primary neurons

After observing robust RSAD2 induction in response to IFN-λ, we next sought to determine how viral infection modulates RSAD2 expression in the presence or absence of the HSV-1 neurovirulence factor ICP34.5 [29]. Immunofluorescence staining analyses of RSAD2 protein in SCGs demonstrated that Δ34.5-infected neurons largely maintained RSAD2 protein levels and perinuclear localization. In contrast, neurons infected with 34.5R showed less punctate organization with RSAD2 dispersed diffusely, including within the neuronal nucleus. This diffusion of RSDA2 may suggest disruption of spatial organization (Figure 7A). The overall RSAD2 protein levels were reduced to non-detectable levels during 34.5R infection particularly at 24 hpi (Figure 7B). Interestingly, RSAD2 gene expression was markedly higher in SCGs infected with the HSV-1 Δ34.5 mutant 24 hpi, reaching levels comparable to those observed during PRV180 infection while the 34.5R showed reduced RSAD2 expression (Figure 7C).

**Figure 7.**
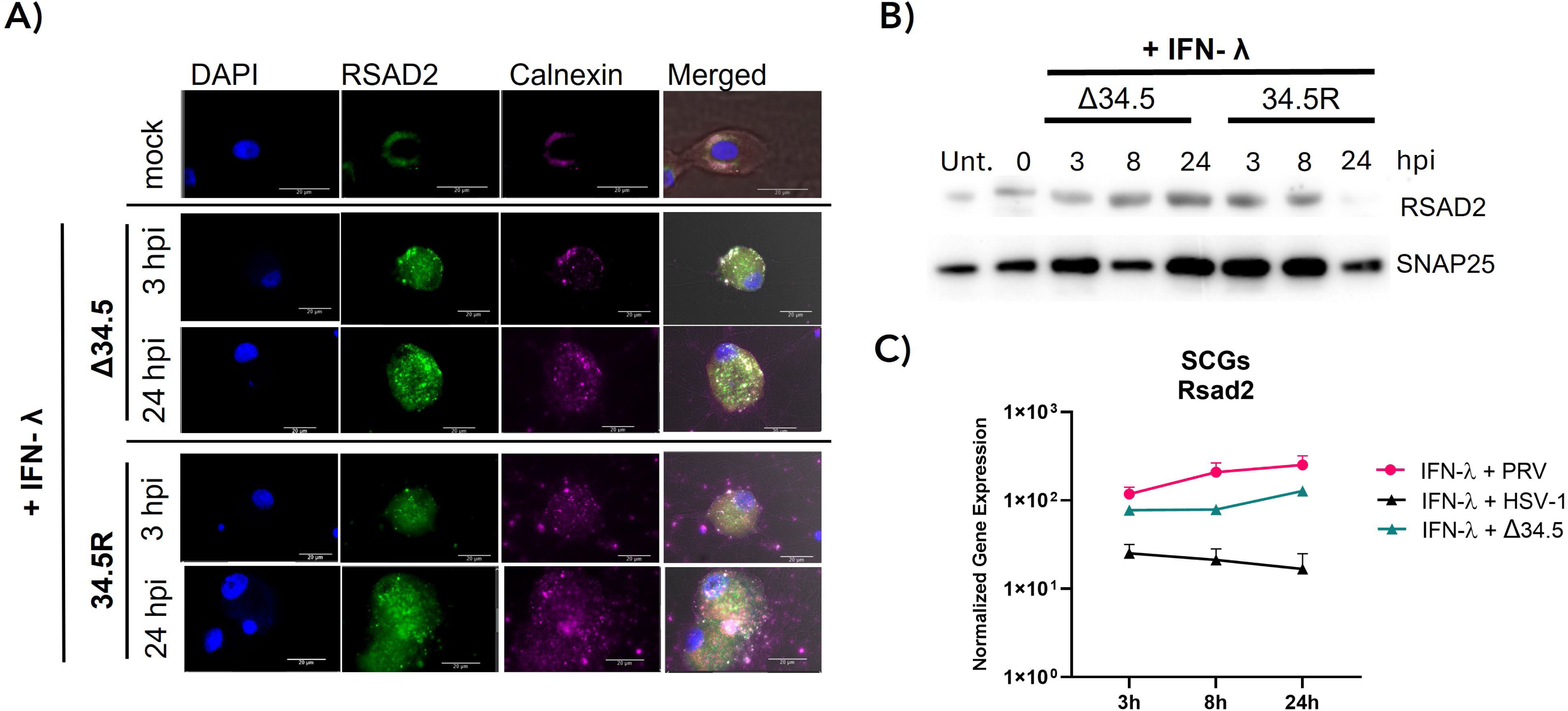
ICP34.5 suppresses RSAD2 expression and disrupts ER localization in primary neurons. (**A**) SCGs were pretreated with rat IFN-λ2 prior to infection with either 34.5R or Δ34.5 at MOI 1. Cells were fixed at 24 hpi and stained with RSAD2 antibody (scale bars, 20 μm). (**B**) Normalized gene expression of RSAD2 in SCGs following infection with Δ34.5, or 34.5R at MOI 1 for 3 h or 24 hpi. Statistical significance was assessed using unpaired two-tailed t tests with Welch’s correction. (**C**) Immunoblotting of SCGs pretreated with rat IFN-λ2 (200 ng/ml) for 24 h and infected with either 34.5R or Δ34.5 at MOI 1 for 3 h, 8 h, or 24 h. Lysates were probed with antibodies against RSAD2 and SNAP25.

### HSV-1 ICP34.5 modulates RSAD2 expression in human neuronal cells

To examine the modulatory roles of ICP34.5 on host response and viral protein production in human neuronal cells, immunofluorescence staining and immunoblotting were used to detect the temporal expression of RSAD2 and viral proteins during Δ34.5 and 34.5R viral infection in IFN-λ-primed SK-N-SH cells. In 34.5R-infected cells, RSAD2 detection decreased at later timepoints of 24 hpi in comparison to earlier timepoints at 3 hpi (Figure 8A). In contrast, Δ34.5-infected cells displayed a more punctate, perinuclear distribution of RSAD2 that persisted at both early and late time points (Figure 8A). In several infected cells, RSAD2 remained concentrated at ER, suggesting that the loss of ICP34.5 preserves its subcellular localization, potentially by preventing viral disruption of host protein trafficking or organelle organization.

**Figure 8.**
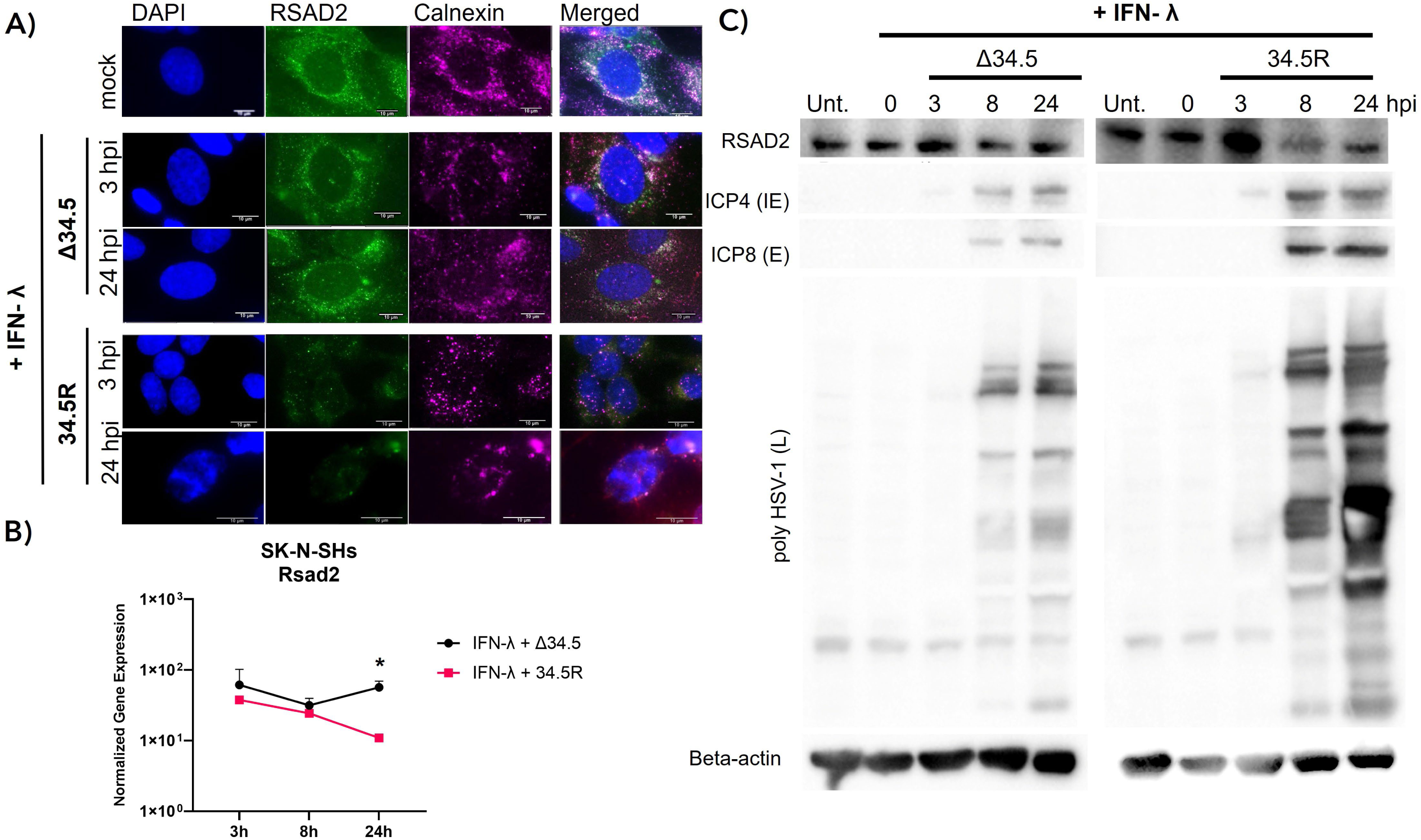
ICP34.5 suppresses RSAD2 expression, disrupts ER localization and supports viral gene expression in SK-N-SH cells. (**A**) Immunofluorescence staining of SK-N-SH cells pretreated with human IFN-λ2 prior to infection with either 34.5R or Δ34.5 at MOI 1. Cells were fixed at 3 hpi and 24 hpi and stained for RSAD2 antibody (scale bars are 10 μm). (**B**) Normalized gene expression of RSAD2 in SK-N-SH cells following infections with either Δ34.5 or 34.5R at MOI 1 across three timepoints: 3, 8, and 24 hpi. Statistical significance was determined using an unpaired Student’s *t* test; *, *p* < 0.05. (**C**) Immunoblotting of SK-N-SH cells pretreated with human IFN-λ2 for 24 h and infected with either 34.5R or Δ34.5 at MOI 1 for 3, 8, and 24 h. Lysates were probed with antibodies against RSAD2, ICP4, ICP8, poly-HSV-1, and beta-actin.

A similar pattern appeared in the steady state expression of transcripts in SK-N-SH cells, where RSAD2 upregulation occurred only after Δ34.5 infection with IFN-λ pretreatment, not with 34.5R virus infection (Figure 8B). These results were confirmed by immunoblot analyses of RSAD2 protein levels. In SK-N-SH cells infected with 34.5R, RSAD2 levels were decreased compared to untreated controls at 24 hpi, whereas Δ34.5-infected cells largely maintained RSAD2 expression (Figure 8C).

To further explore viral infection progress in these cells, we examined HSV-1 viral protein expression. Cells infected with the ICP34.5 revertant strain showed efficient late protein expression at both 8 hpi and 24 hpi, consistent with efficient replication (Figure 8C). In contrast, Δ34.5-infected cells displayed reduced viral protein accumulation (Figure 8C). These results suggest that ICP34.5, by altering the localization and reducing the amounts of RSAD2 protein, promotes HSV-1 protein synthesis in primary rat neurons and human neuronal cells.

### RSAD2 accumulation on the ER sustains p-eIF2α-mediated translational arrest in the absence of functional ICP34.5

To examine whether IFN-λ pretreatment leads to activation of protein synthesis shut off during HSV-1 infection, we assessed eIF2α phosphorylation in SCG neurons infected with the Δ34.5 or revertant strain, 34.5R. SCG neurons pre-treated with IFN-λ and infected with Δ34.5 demonstrated sustained p-eIF2α expression, whereas those infected with 34.5R displayed a gradual reduction in detection levels (Figure 9A).

**Figure 9.**
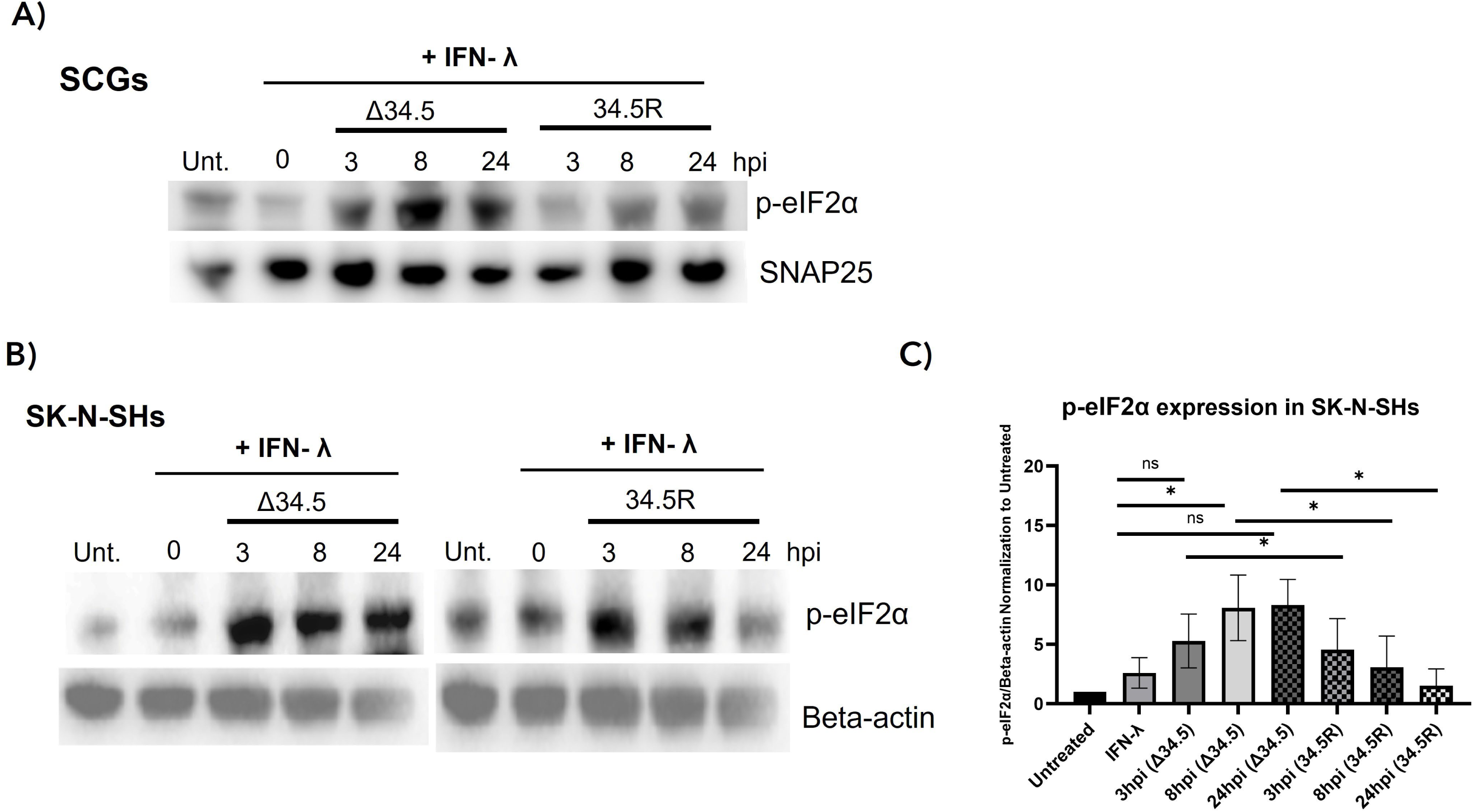
**eIF2α phosphorylation sustains in IFN-λ-treated neurons during Δ34.5 infection.**SCGs were pretreated with rat IFN-λ2 for 24 h and subsequently infected with either the Δ34.5 or 34.5R at MOI 1. (**A**) Cells were harvested 3-, 8-, and 24 h post-infection. Immunoblots were probed with antibodies against phosphorylated eIF2α (p-eIF2α) and Snap25. (**B**) SK-N-SH cells were pretreated with human IFN-λ2 for 24 h, and subsequently infected with Δ34.5 or 34.5R at MOI 1. Cells were harvested 3, 8, and 24 h post-infection for immunoblot analysis. Immunoblots were probed with antibodies against phosphorylated eIF2α (p-eIF2α) and beta-actin. (**C**) RSAD2 levels were normalized to beta-actin and each lane is analyzed for p-eIF2α levels comparing Δ34.5 or 34.5R infection time points (ImageJ, n=3). Statistical significance was determined using an unpaired Student’s *t* test: *, *p* < 0.05, **, *p* < 0.005, ***, *p* < 0.0005, ns, not significant.

SK-N-SH cells infected with the ICP34.5 revertant demonstrated significantly lower levels of p-eIF2α at 8 and 24 hpi, indicating ICP34.5’s ability to recruit protein phosphatase 1 alpha (PP1α) and promote its dephosphorylation [30] (Figure 9B and 9C). In contrast, Δ34.5-infected SK-N-SH cells maintained high levels of eIF2α phosphorylation (p-eIF2α) at all time points, consistent with activation of host stress responses and translational shutdown. These results demonstrate that, in the absence of ICP34.5, IFN-λ-primed neurons maintain high p-eIF2α levels, supporting an antiviral state due to protein translation shut off. These findings reveal that p-eIF2α and RSAD2 serve as key effectors in mediating IFN-λ-induced antiviral restriction, a process that HSV-1 antagonizes through the expression of ICP34.5.

### RSAD2 functions as a key antiviral effector of IFN-λ-mediated response in PNS neurons

To directly assess the role of RSAD2 in mediating IFN-λ-dependent restriction of HSV-1, we performed RNA interference experiments in primary SCG neurons. Immunoblot analysis confirmed efficient silencing of RSAD2 following transfection with a rat RSAD2-specific siRNA compared to a non-targeting (NT) control. Band intensities normalized to SNAP25 demonstrated a significant reduction in RSAD2 protein levels, validating the knockdown efficiency (Figure 10A).

**Figure 10.**
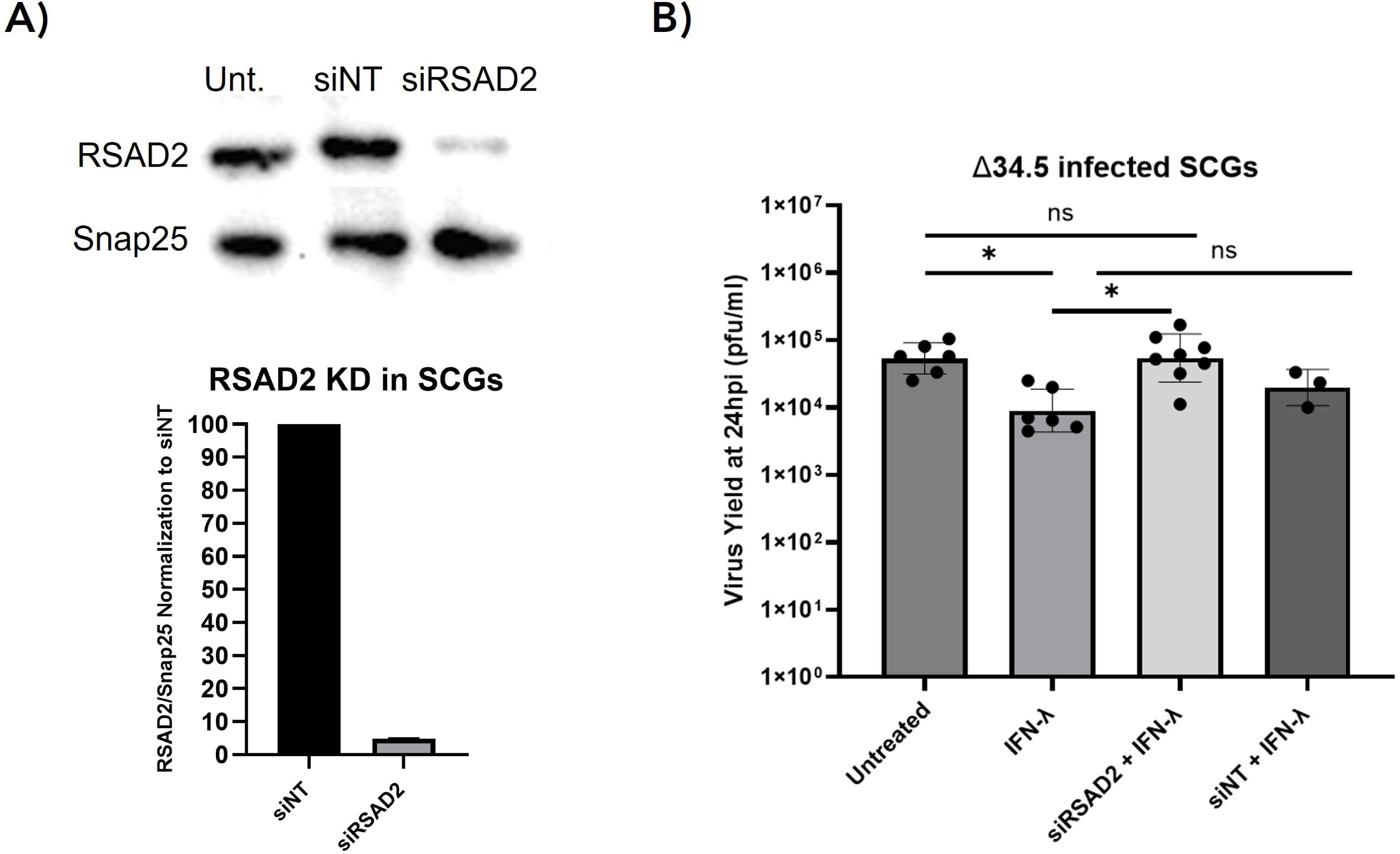
RSAD2 knockdown restores ICP34.5-deficient HSV-1 replication in IFN-λ-treated SCGs. (**A**) Representative immunoblot showing RSAD2 protein levels in SCGs following transfection with nontargeting (NT) or RSAD2-specific siRNA. Quantification of the bands shown below the blot using ImageJ. RSAD2 levels were normalized to SNAP25 and demonstrate efficient RSAD2 depletion (RSAD2, 0.061 ± 0.003 SEM relative to NT control; unpaired Student’s *t* test, **, *p* < 0.005). (**B**) SCGs were transfected with siNT or]siRSAD2 for 48 h and pretreated with rat IFN-λ2 for 24 h prior to infection with the HSV-1 Δ34.5 at an MOI of 10. Viral titers were quantified at 24 h post-infection. Bars represent mean ± SEM. Individual points indicate biological replicates. Statistical significance was determined using Brown-Forsythe and Welch one-way ANOVA followed by Games-Howell’s multiple comparisons test. *, p < 0.05; ns, not significant.

We next evaluated the functional impact of RSAD2 depletion on viral replication during IFN-λ treatment. SCGs were pretreated with rat IFN-λ and subsequently infected with the HSV-1 Δ34.5 mutant for 24 h at MOI of 10. In siNT-transfected neurons, IFN-λ pretreatment significantly restricted Δ34.5 replication, consistent with earlier observations. In contrast, RSAD2 knockdown partially rescued viral replication, resulting in a significant increase in viral titers compared to the siNT controls (p < 0.05; Figure 10B). These findings demonstrate that RSAD2 contributes to the antiviral effect of IFN-λ in neurons and plays a central role in restricting HSV-1 infection in the absence of ICP34.5.

## Discussion

Neurons occupy a uniquely vulnerable position in antiviral immunity, requiring defense strategies that restrict viral replication without compromising long-term cellular viability. Building on our previous findings, this study further characterizes IFN-λ responses in peripheral neurons and identifies RSAD2 (viperin) as a key effector mediating antiviral responses against α-HV and demonstrates that HSV-1 has evolved unique strategies to mitigate these defenses. We employed an RNA-sequencing-based approach to comprehensively characterize IFN-λ-mediated responses in neurons, identifying strong induction of RSAD2 and other ISGs such as Mx1 and USP18. Time-course analysis revealed distinct kinetic differences in neurons treated with either type I or type III IFNs. Type I IFN treatment resulted in a rapid, high-magnitude induction of ISG expression, while IFN-λ induced a slower, lower magnitude of induction. This pattern, previously observed in polarized epithelial cells, appears more pronounced in neurons [31].

Neurons are seemingly predisposed to a low but constant response when stimulated by IFN-λ. The variations in receptor levels (IFN-λR1/IL-10R2), the speed of signal transduction, or the differential accessibility of chromatin regions for ISGs could all play roles in this reduced sensitivity. The delayed yet prolonged response to IFN-λ might be advantageous for neurons, as an overt or rapid activation of antiviral pathways could cause cytotoxic stress and damage. These findings are supported by studies demonstrating that neurons exhibit distinct responses to inflammatory cytokines compared to proliferating cells, likely reflecting adaptations to preserve neuronal integrity and function [32, 33].

Notably, RSAD2 was the most upregulated ISG in IFN-λ-treated neurons. Although its role in epithelial cells is well characterized, RSAD2’s role and function in neurons remain to be elucidated [34, 35]. Our results demonstrate that RSAD2 is strongly induced by IFN-λ in peripheral neurons and localizes to punctate perinuclear ER-associated structures [36]. RSAD2’s subcellular localization is functionally critical for its antiviral activity. The protein predominantly localizes to membranes derived from the ER, lipid droplets, and mitochondria through its N-terminal amphipathic α-helix [36]. This membrane association positions RSAD2 at advantageous cellular sites frequently used by viruses for replication and assembly. For α-HVs, which undergo secondary envelopment at trans-Golgi network to acquire their final lipid envelope, RSAD2’s strategic localization positions it to interfere with late stages of the viral lifecycle.

IFN-λ-induced RSAD2 accumulation in human SK-N-SH cells, with prominent perinuclear puncta clearly colocalizing with the ER marker calnexin by immunofluorescence. Likewise, rat primary SCG neurons exhibited similar induction; albeit with a more dispersed punctate staining matching the localization of ER. Peripheral neurons possess specialized ER domains and organization adapted to their unique morphology, including extensive axonal and dendritic processes, which may make RSAD2 localization particularly critical where the ER structure is different from epithelial cells [37–39].

When we examined the potency of IFN-λ-mediated antiviral response against α-HVs, the results were striking. While PRV remained highly sensitive to IFN-λ restriction in neurons, HSV-1 replicated efficiently despite IFN-λ inducing comparable STAT1 phosphorylation and ISG expression [9, 40]. This disparity demonstrated the fundamental difference in how these two α-HVs interact with neuron-specific immune defenses. Specifically, HSV-1’s resistance to IFN-λ is neuron-specific as opposed to a general phenotype. Both rat and human fibroblasts displayed significant IFN-λ-mediated viral suppression during HSV-1 infection, demonstrating that fibroblasts retain the capacity to restrict HSV-1 infection. Additionally, this neuron-specific resistance is conserved across species as both rat SCGs and human SK-N-SH neuroblastoma cells failed to restrict HSV-1, ruling out species-specific differences. These findings indicate that HSV-1 has evolved specialized mechanisms to overcome neuron-specific antiviral defenses

The absence of ICP34.5 in PRV and its presence in simplexviruses (HSV-1 and HSV-2) provided a natural genetic tool to dissect its role in overcoming neuronal IFN-λ responses, while highlighting differences in α-HVs neuroinvasion. Infection with an ICP34.5-deficient HSV-1 mutant (Δ34.5) restored sensitivity to neuronal IFN-λ pretreatment, demonstrating that ICP34.5 is necessary for HSV-1 to overcome neuron-specific antiviral defenses. Importantly, Δ34.5 replicated to yield wild-type levels of progeny in untreated neurons, confirming that its IFN-λ sensitivity reflects loss of immune evasion capacity rather than baseline replication defects. The role of ICP34.5 in overcoming IFN-λ defenses highlights key differences in α-HV neuroinvasion strategies.

The molecular basis for ICP34.5’s ability to overcome neuronal IFN-λ-RSAD2 mediated defenses involves complementary mechanisms that collectively disable host antiviral responses blocking protein synthesis. We propose that ICP34.5 first disrupts RSAD2’s functional spatial organization. In IFN-λ-primed neurons infected with wild-type HSV-1, RSAD2 puncta rapidly dispersed from perinuclear foci to diffuse distribution in the cytoplasm, coinciding with a reduction in total RSAD2 protein levels. In contrast, Δ34.5 infection preserved RSAD2 localization specifically at the ER. This phenotype correlates with strong viral restriction, supporting the model that proper RSAD2 localization is crucial for its antiviral role.

Secondly, ICP34.5 reverses RSAD2’s downstream effects on translational shut off. ICP34.5 recruits protein phosphatase 1α (PP1α) to dephosphorylate eIF2α, thereby reversing the translational shutoff imposed by protein kinase R (PKR) and other stress-activated eIF2α kinases during viral infection [30, 41]. The protein kinase R-like endoplasmic reticulum kinase (PERK) pathway activates the unfolded protein response (UPR) via inhibition of translation, reducing ER load during stress. During ER stress, PERK phosphorylates eIF2α, thereby activating ATF4-mediated stress responses and cell arrest. This function is critical because eIF2α phosphorylation normally blocks both viral and cellular protein synthesis as a protective antiviral mechanism [42].

By restoring translation, ICP34.5 enables continued viral protein synthesis despite PKR activation and may allow viral proteins to antagonize ISG effector functions.

Our immunoblot analysis confirmed this mechanism across both neuron types, demonstrating that Δ34.5-infected SK-N-SH cells maintained high levels of phosphorylated eIF2α and exhibited severely impaired late viral protein synthesis. Similarly, immunoblot analysis of IFN-λ-pretreated rat SCGs infected with Δ34.5 or 34.5R demonstrated that viral protein detection was reduced in Δ34.5 infection, consistent with human SK-N-SH cell results. In contrast, 34.5R infections showed eIF2α dephosphorylation and late protein production in both cell types, demonstrating that ICP34.5 antagonizes eIF2α-mediated translational arrest. The consistency of this phenotype across primary rat neurons and human immortalized neuronal cell lines demonstrates that ICP34.5’s ability to reverse eIF2a phosphorylation and restore viral translation is a fundamental mechanism conserved across neuronal cells.

Additional ICP34.5 functions complement these mechanisms. ICP34.5 inhibits autophagy by binding Beclin-1, preventing autophagosome formation and the subsequent degradation of viral components [19, 20]. Manivanh et al. demonstrated that ICP34.5 blocks TANK-binding kinase 1 (TBK1)-mediated phosphorylation of interferon regulatory factor 3 (IRF3), thereby attenuating type I interferon production in trigeminal neurons (TG) [18]. By targeting RSAD2 localization, eIF2α-mediated translational control, autophagy, and IRF3-dependent interferon induction, ICP34.5 functions as a key antagonist that counteracts neuronal antiviral defenses.

To confirm RSAD2’s role in restricting HSV-1 lacking ICP34.5, we performed loss-of-function studies in neurons. siRNA-mediated RSAD2 knockdown in IFN-λ-primed SCG neurons partially restored replication of Δ34.5 HSV-1, indicating that RSAD2 loss is only consequential when the virus lacks ICP34.5. This strongly suggests that RSAD2 and ICP34.5 function as opposing determinants of infection outcome in neurons. In Δ34.5 infection, RSAD2 remains localized in ER associated compartments, maintains high mRNA and protein levels, and potently restricts infection by interfering with viral protein synthesis. ICP34.5 counteracts the antiviral responses enabling the virus to proceed with efficient gene expression despite the IFN-λ primed antiviral state. This two-pronged antagonism, spatial disruption of RSAD2 plus functional reversal of its downstream effects on translation, demonstrates the selective pressure RSAD2 imposes on HSV-1 and explains why viruses lacking these countermeasures (e.g., PRV) remain vulnerable to neuronal IFN-λ responses.

HSV-1 uses multifaceted strategies to suppress RSAD2, highlighting its relevance as a viral restriction factor and suggesting evolutionary selection pressure. The HSV-1 virion host shutoff protein (vhs), encoded by UL41, is a multifunctional endoribonuclease that nonspecifically degrades host mRNAs to shut off cellular protein synthesis and redirect the translational machinery toward viral proteins. Studies have demonstrated vhs targets RSAD2 mRNA and deplete RSAD2 protein during infection. Shen et al. demonstrated that RSAD2 strongly inhibits UL41-null HSV-1 replication, supporting the idea that the virus actively suppresses RSAD2 [43]. However, in IFN-λ-primed neurons where RSAD2 protein is pre-induced and accumulated before viral entry, vhs-mediated mRNA degradation may be insufficient to eliminate pre-existing RSAD2 protein, which has a relatively long half-life (about 6-9 hours in many cell types) [44]. Under these conditions, ICP34.5 provides a complementary and perhaps more critical mechanism by disrupting RSAD2 function at the protein level. The existence of two independent viral mechanisms targeting RSAD2 underscores its importance as an antiviral restriction factor and reveals an evolutionary arms race between host and pathogen.

The lack of an ICP34.5 homologue in PRV, which would otherwise enable direct assessment of RSAD2 function in the absence of viral impact, provides a critical comparison. This study demonstrated that PRV infection maintained RSAD2 expression during infection at 3 hpi, 8 hpi, and 24 hpi when neurons were primed with IFN-λ for 24 h. The parallels between PRV and Δ34.5 HSV-1 sensitivity strongly suggest that PRV infection fails to fully overcome RSAD2-mediated defenses. This finding has critical implications for understanding α-HV evolution and neurotropism. PRV, despite being a highly neurotropic swine virus, does not naturally infect higher primates. VZV, a human varicellovirus which also lacks ICP34.5, establishes latency predominantly in human dorsal root ganglia but requires higher viral loads and demonstrates limited spread in PNS neurons [45–47]. The absence of ICP34.5 in varicelloviruses may render them more vulnerable to IFN-λ-mediated responses, partially explaining their distinct neurobiological properties compared to simplexviruses [48]. The differential sensitivity of PRV (restricted) versus HSV-1 (resistant) to IFN-λ represents that RSAD2 and other ISG restriction are potent and effective when viral countermeasures are absent, but can be overcome by viruses encoding specialized antagonists such as ICP34.5. The role of ICP34.5 in overcoming IFN-λ defenses highlights fundamental differences in α-HV neuroinvasion strategies and may explain differential capacities for latency infection maintenance and reactivation [49].

Based on the findings in this study, we propose a model for how IFN-λ establishes antiviral response in neurons and how HSV-1 overcomes them via ICP34.5 antagonism (Figure 11). In this model, IFN-λ produced by infected epithelial cells at mucosal barriers primes uninfected cells, including PNS axon terminals innervating these sites. IFN-λ binding to receptors, IFNLR1/IL10B, activates STAT1-selective signaling cascades, leading to transcription of ISGs including RSAD2. RSAD2 protein accumulates at ER-derived membranes [36]. Upon HSV-1 entry and transcription initiation of early transcription events, PKR is activated by viral dsRNA, leading to the phosphorylation of eIF2α which results in translational arrest and induction of autophagy [19]. ICP34.5’s role in mitigating PKR-mediated translational shutoff through recruitment of PP1α is well established [30, 41]. Our findings further suggest that RSAD2 imposes an additional antiviral defense in peripheral neurons by inducing ER stress, which activates PERK, resulting in expression of the unfolded protein response (UPR) pathway [50]. This RSAD2-mediated stress response acts as a second translational checkpoint that reinforces the antiviral state in peripheral neurons. However, ICP34.5 antagonizes both PKR and PERK pathways by dephosphorylating eIF2α, limiting RSAD2’s role in antiviral defense and enabling HSV-1 to establish production infection in neurons despite IFN-λ priming.

**Figure 11.**
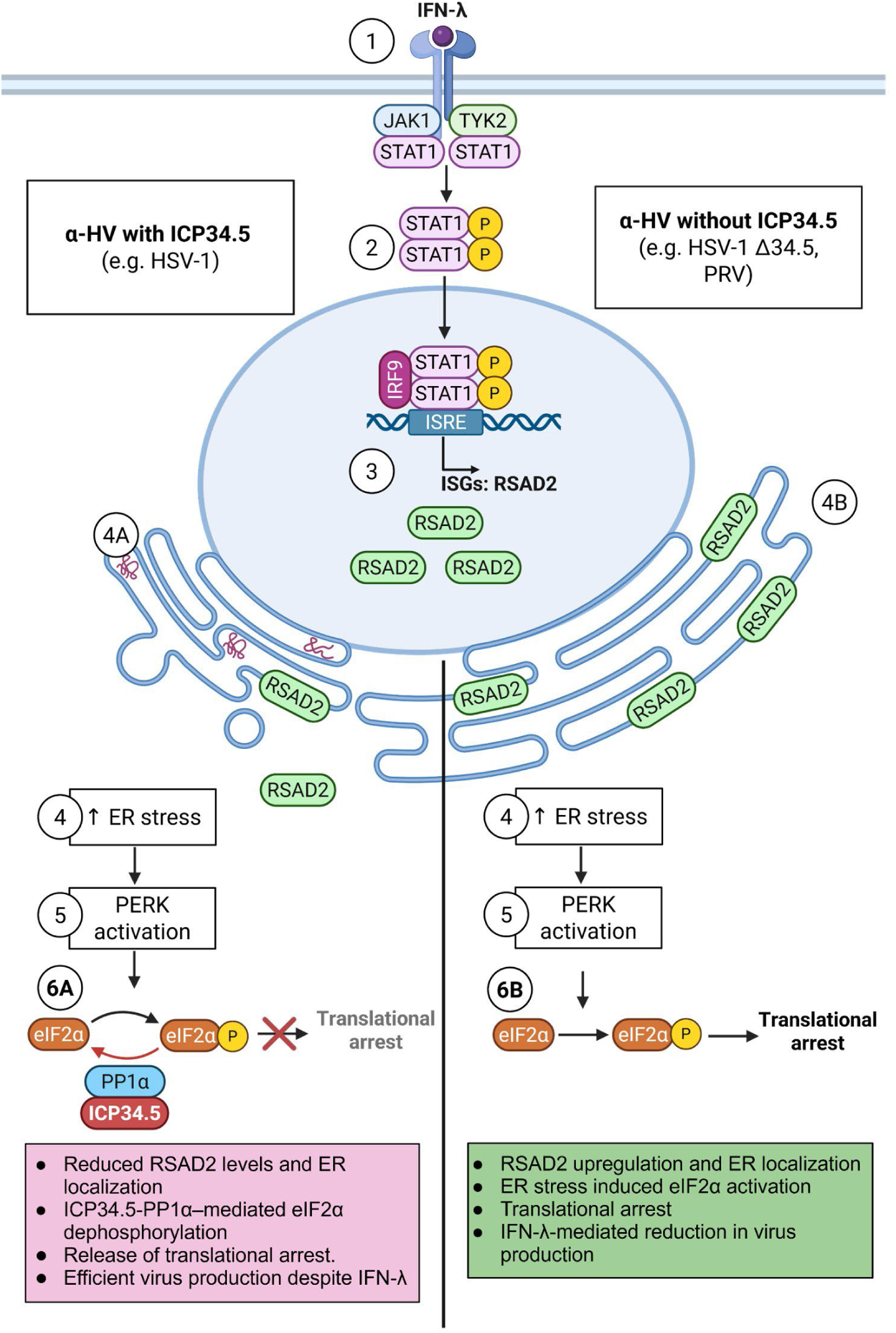
Model of ICP34.5-mediated regulation of RSAD2 during IFN-λ-induced antiviral responses in neurons. IFN-λ binding to its receptor initiates the intrinsic immune response (**1**), inducing the phosphorylation of JAK1/TYK2 and formation of STAT1 homodimers in peripheral neurons (**2**), which promote the expression of a subset of interferon stimulated genes (ISGs), including RSAD2 (**3**). RSAD2 localizes to the endoplasmic reticulum (ER) (**4**). RSAD2-mediated ER stress activates the PKR-like endoplasmic reticulum kinase (PERK) to eIF2α and translational arrest (**5**). This leads to ATF4 nuclear localization and induction of the unfolded protein response (UPR) (**6A**). ICP34.5 counteracts this antiviral response by recruiting PP1α, which dephosphorylates eIF2α, reversing translational shut off (**6B**).

When HSV-1 infects IFN-λ primed neurons (via axon termini or cell bodies), multiple antiviral mechanisms are triggered. RSAD2 activity and associated ER stress are consistent with activation of PERK and PKR, leading to eIF2α phosphorylation that enforces translational arrest. HSV-1 overcomes this barrier by deploying ICP34.5 to dismantle RSAD2’s structural organization while simultaneously subverting the integrated stress response to restore viral translation. In contrast, viruses lacking ICP34.5 cannot disrupt RSAD2 localization or reverse eIF2α phosphorylation, resulting in sustained translational arrest and inefficient infection. Together, RSAD2 and ICP34.5 define a neuron-specific antagonistic mechanism that determines the efficiency of HSV-1 infections in a type III IFN-primed environment. This model establishes a framework for understanding how different α-HVs have evolved distinct strategies to navigate the neuronal innate immune landscape.

## Materials and Methods

### Cells and Viruses

Rat2 cells were purchased from ATCC. The cells were cultured in Dulbecco Modified Eagle medium (DMEM) (Cytiva parent company Danaher Corporation, Washington, DC, USA), 1% penicillin-streptomycin (PS) (Cytiva parent company Danaher Corporation, Washington, DC, USA), and 10% fetal bovine serum (FBS) (Genessee, Burlington, MA, USA). Human dermal fibroblasts (HDFs) were provided by Nir Drayman Laboratory (University of California, Irvine) and grown in DMEM, 1% PS, and 10% FBS. Porcine kidney epithelial cell line (PK15) was grown in DMEM, 1% PS, and 5% FBS. Human neuroblastoma cell line, SK-N-SH cells were generously shared by Bert Semler Laboratory (University of California, Irvine) and grown in DMEM, 1% PS, and 20% FBS. PRV-Becker is a wild-type laboratory strain [52]. PRV-180 expresses mRFP1-VP26 in a PRV-Becker background [27]. HSV-1 OK14, which expresses an mRFP-VP26 capsid tag, was previously described [26]. HSV-1 Δ34.5 and 34.5R were obtained from Nir Drayman (University of California, Irvine) with the permission of David Leib (Dartmouth Geisel School of Medicine). Δ34.5, originally known as *17terma*, has been previously described along with its revertant, 34.5R, respectively [51, 53]. All HSV-1 or PRV strains were propagated and titered by plaque assay on Vero or PK15 cells, respectively, in DMEM supplemented with 2% FBS and 1% penicillin-streptomycin.

### Primary Neuronal Cultures

Superior cervical ganglionic (SCG) neurons were used for all primary neuron experiments that were cultured in modified Campenot tri-chambers on 35-mm or 12-well dishes (Corning, Corning, NR, USA). Dishes were coated with poly-DL-ornithine (Millipore Sigma, Burlington, MA, USA) and mouse laminin (Life Technologies Carlsbad, CA, USA) to promote neuronal adherence. SCG were isolated from 16- or 17-day Sprague Dawley rat embryos (Charles River Laboratories), as described by Curanovic et al. [54]. Neurons were cultured in neurobasal medium (Gibco, Billings, MT, USA) + 50X B-27 supplement (Gibco, Billings, MT, USA) + 100X Penicillin-Streptomycin-Glutamine (Gibco, Billings, MT, USA) + 1000X murine nerve growth factor (Gibco, Billings, MT, USA). Two days after seeding, 1 μM Cytosine β-D-arabinofuranoside (Ara-C, Millipore Sigma, Burlington, MA, USA) was added to select against any mitotic cells. Neuronal medium was changed every five to seven days. All animal work was performed in accordance with the Institutional Animal Care and Use Committee of the University of California, Irvine Research Board under protocol: AUP-24-008.

### Immunofluorescence staining

Primary SCG neurons were cultured on poly-D-lysine and laminin-coated glass coverslips and processed for immunofluorescence following the experimental treatments described in each figure. Neurons and SK-N-SH cells were fixed in 4% paraformaldehyde (Electron Microscopy Sciences, Hatfield, PA, USA) for 15 minutes at room temperature (RT), washed three times in PBS, permeabilized with methanol for 5 minutes at-20C, and blocked in 5% BSA in PBS for 1 hour. Primary antibodies were diluted in blocking buffer and incubated overnight at 4°C, including anti-RSAD2 (Invitrogen, Waltham, MA, USA; 1:500) and anti-calnexin (Santa Cruz Biotechnology, Santa Cruz, CA, USA; 1:250). Following primary incubation, cells were washed in PBS and incubated with Alexa Fluor-conjugated goat anti-mouse or goat anti-rabbit secondary antibodies (Thermo Fisher Scientific, Waltham, MA, USA; 1:500) for 1 hour at RT. Nuclei were stained with DAPI (1 µg/mL) for 1 hour at RT, along with secondary antibodies at 1:1000, and coverslips were mounted using Polygel mountant (Thermo Fisher Scientific). Z-stacked images were acquired using a Leica DMI8 microscope with 20X or 63X oil-immersion objectives under consistent settings for each comparison. Image analysis was performed using standardized thresholding and background subtraction to ensure uniform exposure across samples using Leica Imaging Software.

### Interferon treatment

Recombinant rat IFN-λ2 (200 ng/mL) was added directly to neuronal cultures or fibroblasts for 24 hours prior to infection (VWR, Radnor, PA, USA). SK-N-SH cells were treated with human IFN-λ2 (200 ng/mL, 24 hours) (Thermo Fisher, Waltham, MA, USA).

### Real-Time Quantitative PCR

Viral transcription was quantified using primer sets designed for immediate-early, early, and late α-herpesvirus genes. Primers were obtained from Integrated DNA Technologies (IDT, Coralville, IA, USA) (Table 1). RNA isolation, cDNA synthesis, and qPCR amplification were performed as described above, and viral transcript abundance was calculated using the comparative Ct method (2−ΔΔCt) normalized to GAPDH.

**Table 1.**
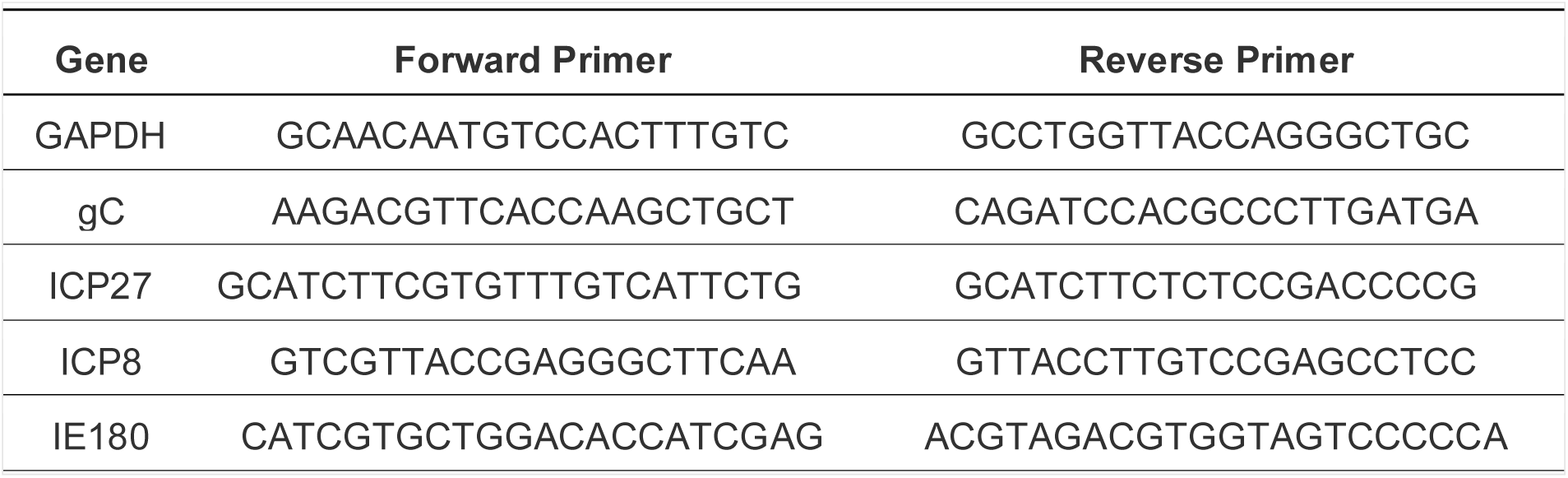
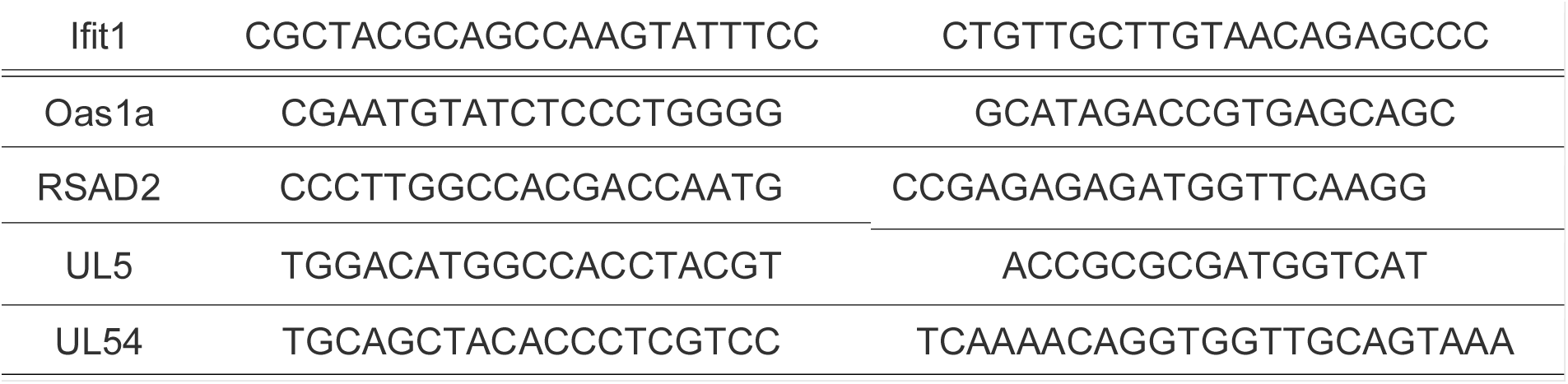
Primer sequences used in this study.

### RNA Isolation and RNA-Sequencing

Total RNA for RNA-sequencing was isolated from SCG neurons using the RNeasy Mini Kit (Qiagen, Hilden, Germany), following the manufacturer’s column-based protocol to ensure high-purity RNA suitable for downstream library preparation. SCGs from 2 wells of a 12-well plate were lysed directly in RLT buffer supplemented with β-mercaptoethanol. Lysates were homogenized using spin columns (Qiagen) prior to column binding. Genomic DNA contamination was removed using on-column RNase-free DNase I digestion (Qiagen) according to the manufacturer’s guidelines. Samples were washed and eluted in nuclease-free water, snap-frozen, and stored at-20°C until quality assessment. Only samples with RNA integrity numbers (RIN) suitable for library preparation were submitted for sequencing. All RNA isolations were performed in RNase-free conditions, and all neuronal samples were processed in parallel to minimize batch effects during downstream transcriptomic analyses. Polyadenylated mRNA was enriched using the NEBNext Poly(A) mRNA Magnetic Isolation Module following the manufacturer’s protocol (New England Biolabs, Ipswich, MA). Strand-specific RNA-seq libraries were constructed utilizing the NEBNext Ultra II RNA Library Prep Kit with Sample Preparation Beads using manufacturer instructions (New England Biolabs). Library amplification was performed by PCR (8-12 cycles, optimized per input RNA quality) and barcoded using the NEBNext Multiplex Oligos for Illumina – 96 unique Dual Index Primers (New England Biolabs). Libraries were normalized and pooled prior to submission for Illumina paired-end sequencing (2×150 bp NovaSeq 6000) by the UCI Genomics core. Raw sequencing reads were evaluated for quality using FastQC. Differential gene expression analysis was conducted with DESeq2 in RStudio. Genes with an adjusted p-value < 0.05 and log₂ fold change > 1 were deemed significantly differentially expressed. Gene ontology (GO) and pathway enrichment analyses were carried out using gProfiler [55].

### siRNA Knockdown of RSAD2 in SCG Neurons

RSAD2 knockdown was performed using rat RSAD2-specific siRNA delivered via siMagnetofection, optimized for primary neuron transfection (OzBiosciences, San Diego, CA, USA). Neurons were treated with either RSAD2 siRNA or a non-targeting (NT) control siRNA according to the manufacturer’s instructions at 50 nM (Horizon Discovery, Cambridge, UK). After 48 hours, knockdown efficiency was validated via immunoblotting. Protein lysates were collected in RIPA buffer supplemented with protease and phosphatase inhibitors, quantified using a BCA assay, and resolved by SDS-PAGE. Membranes were probed with antibodies against anti-RSAD2 (1:5000) (Invitrogen, Waltham, MA, USA) and SNAP25 (1:2000) (loading control) (Cell Signaling Technology, Danvers, MA, USA). Band intensities were quantified in *ImageJ* and normalized to SNAP25. SCGs were pretreated with IFN-λ2 for 24 hours, then infected with HSV-1 Δ34.5 at a multiplicity of infection (MOI) of 10 for 24 hours prior to harvest.

## Statistical Analysis

All graphical analyses were conducted in *GraphPad Prism*. An unpaired Student’s *t* test was used as indicated. Additionally, one-way analysis of variance (ANOVA) was applied, where variance homogeneity could not be assumed; Brown-Forsythe and Welch ANOVA tests were used. Upon significance, Games-Howell post hoc tests were applied for multiple comparisons. Data are presented as mean ± SEM. Statistical significance thresholds were defined as follows: ns, not significant; *, p < 0.05; **, p < 0.005, ***, p < 0.0005; ****, p < 0.00005.

### Western Blot Analysis

Cell lysates were prepared using RIPA-light buffer including 1 mM Dithiothreitol (DTT) and a protease inhibitor cocktail (Roche Holding AG, Basel, Switzerland). Following 30 min incubation on ice and sonication, cells were pelleted, and the supernatant was placed in a new tube with 5X Laemmli buffer before boiling at 95 °C. Immunoblotting was performed using anti-Beta-actin (1:10,000) (Millipore Sigma, Burlington, MA, USA), anti-p-eIF2α (1:2000) (Cell Signaling Technology, Danvers, MA, USA), anti-SNAP25 (1:2000) (Cell Signaling Technology, Danvers, MA, USA), anti-RSAD2 (Invitrogen, Waltham, MA, USA), anti-ICP4 (Abcam, Cambridge, UK), anti-ICP8 (Abcam, Cambridge, UK), anti-gC (Abcam, Cambridge, UK), and anti-HSV-1 (1:1000) (Agilent Dako). Secondary mouse or rabbit antibodies (Millipore Sigma, Burlington, MA, USA, 1:5000) were used following primary antibody incubation.

## Acknowledgements

This work was supported in part by NIH grants R01AI185349 (O.O.K.), NIH T32 fellowships AI007319 (S.S.), AG081185 (K.T.Y.L), NS121727 (A.L.). We thank Dr. David Leib (Dartmouth Geisel School of Medicine) and Dr. Bert Semler (University of California, Irvine) for generously sharing reagents.

## Data Availability

Raw RNAseq data were deposited at the National Center for Biotechnology Information (NCBI) Sequence Read Archive (SRA) with an accession number: PRJNA1465816. https://urldefense.com/v3/ https://dataview.ncbi.nlm.nih.gov/object/PRJNA1465816?reviewer= nkaip9ak89e7ebph3fhiov1tnc;!!CzAuKJ42GuquVTTmVmPViYEvSg!JsS72vWP5OpGdjnL2ck baBSwvR78XbpV-2u9GT5oZSiPmJlksmWU55NeEIHTAGzrvUYtrrfF00k_wtk4$

## Supplemental Information

**Supplemental Figure 1.**
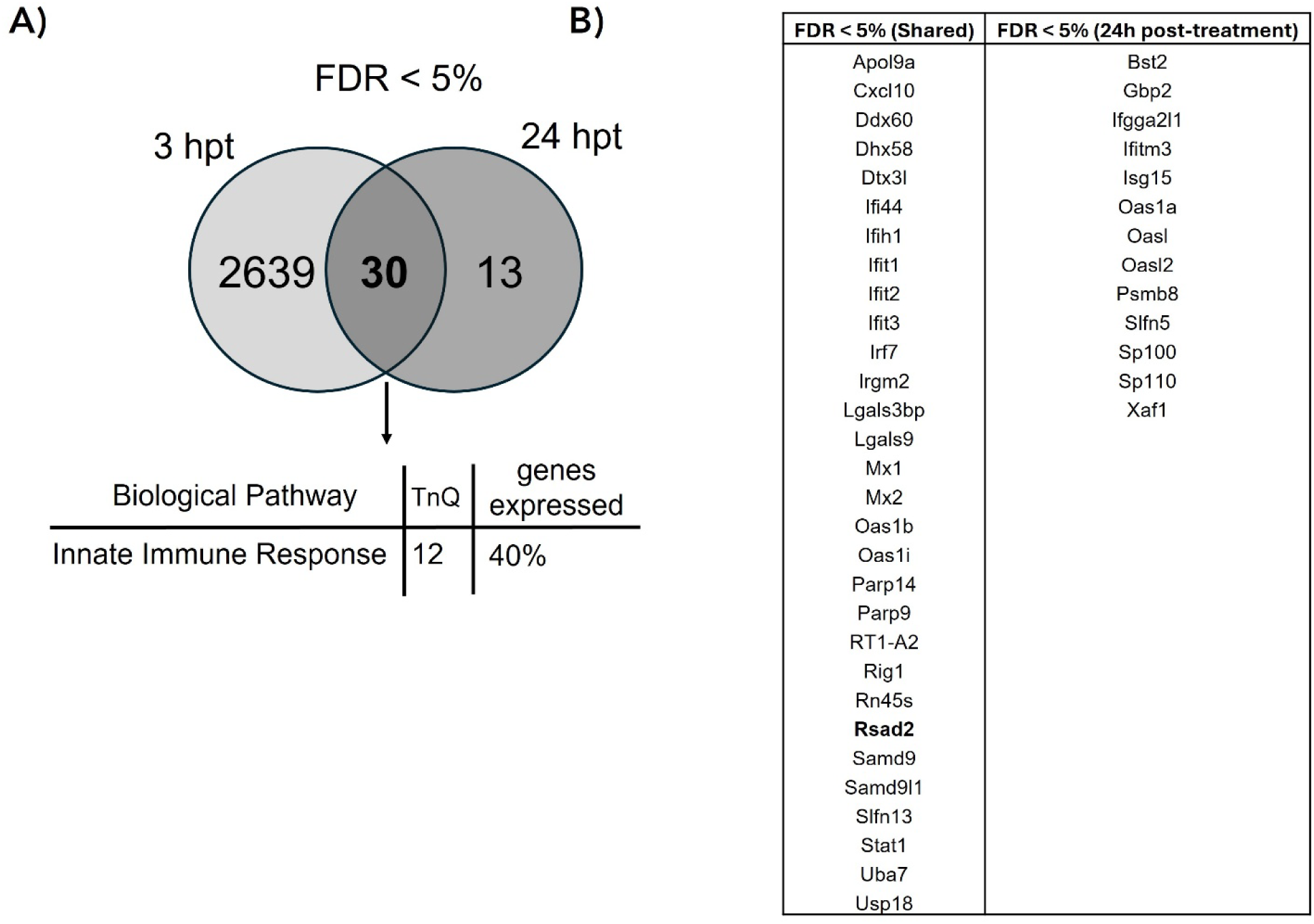
Differential gene expression profiles at early and late timepoints following IFN-λ treatment in SCGs. (A) Venn diagram of differentially expressed genes (DEGs) (FDR <5%) comparing early (3 h) and late (24 h) responses. Shared genes are primarily linked to the innate immune response (40%). *gProfiler* was utilized to categorize genes. *GeneVenn* was used to identify genes shared after 3 and 24 hpt with IFN-λ. (B) Table demonstrating significantly induced genes (FDR <5%) shared between 3 h and 24 hpt primary neurons (left) and uniquely enriched genes at 24 hpt (right).

**Supplemental Figure 2.**
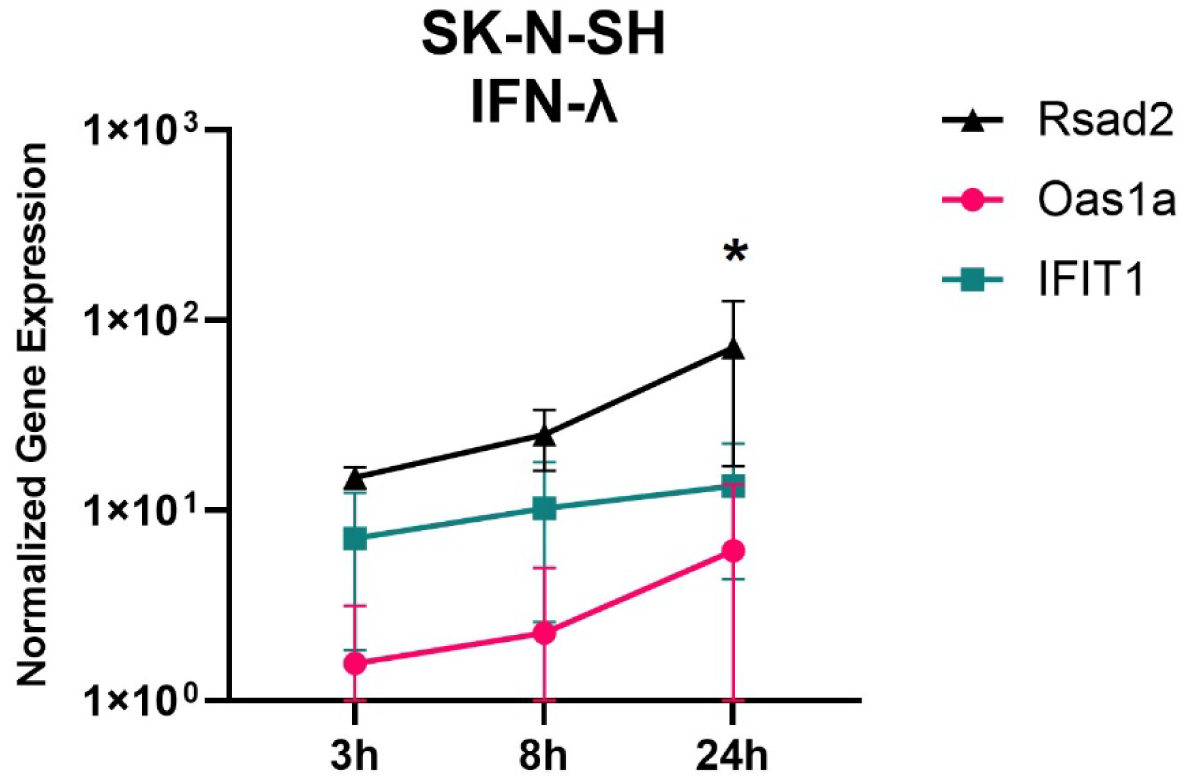
ISG expression kinetics in SK-N-SH cells. RSAD2, Oas1a and IFIT1 expressions were quantified by Q-PCR relative to GAPDH following human IFN-λ treatment for 3-, 8- and 24-hours. Statistical significance was performed using Student’s *t* test: *, *p* < 0.05 (n=3 replicates).

